# GrapeTree: Visualization of core genomic relationships among 100,000 bacterial pathogens

**DOI:** 10.1101/216788

**Authors:** Zhemin Zhou, Nabil-Fareed Alikhan, Martin J. Sergeant, Nina Luhmann, Cátia Vaz, Alexandre P. Francisco, João André Carriço, Mark Achtman

## Abstract

1. Current methods struggle to reconstruct and visualise the genomic relationships of ≥100,000 bacterial genomes.
2. GrapeTree facilitates the analyses of allelic profiles from 10,000’s of core genomes within a web browser window.
3. GrapeTree implements a novel minimum spanning tree algorithm to reconstruct genetic relationships despite missing data together with a static “GrapeTree Layout” algorithm to render interactive visualisations of large trees.
4. GrapeTree is a stand-along package for investigating Newick trees plus associated metadata and is also integrated into EnteroBase to facilitate cutting edge navigation of genomic relationships among >160,000 genomes from bacterial pathogens.
5. The GrapeTree package was released under the GPL v3.0 Licence.

## Introduction

Twenty years ago, MultiLocus Sequence Typing (MLST) was introduced to elucidate and characterise the population structure of bacterial pathogens (Maiden et al., 1998). MLST schemes were rapidly implemented for multiple bacterial species, usually on the basis of sequences of seven housekeeping gene fragments (loci), whose unique sequences (alleles) were each assigned a unique integer number (Jolley & Maiden, 2014). The ordered combination of allelic integers forms the ST (Sequence Type). We refer to this form of MLST as legacy MLST.

Legacy MLST lacks the resolution needed for epidemiological tracking of transmission networks and disease control, and recent attention has focussed on higher resolution MLST schemes based on the entire genome (whole genome MLST-wgMLST) (Nadon et al., 2017), or on the core genes that are present in most isolates of a species or genus (core genome MLST-cgMLST) (Maiden et al., 2013). Various such schemes have now been described (Table 1). wgMLST or cgMLST data are visualised and/or analysed with methods which were quite effective for legacy MLST, including minimum spanning trees (goeBURST/SeqSphere+/Bionumerics), phylograms (UPGMA or NJ [Neighbor-Joining]) and NeighborNet (SplitsTree4). However, cgMLST and wgMLST data present novel problems that did not arise with legacy MLST due to the big data conundrum and missing data.

**Table 1.**
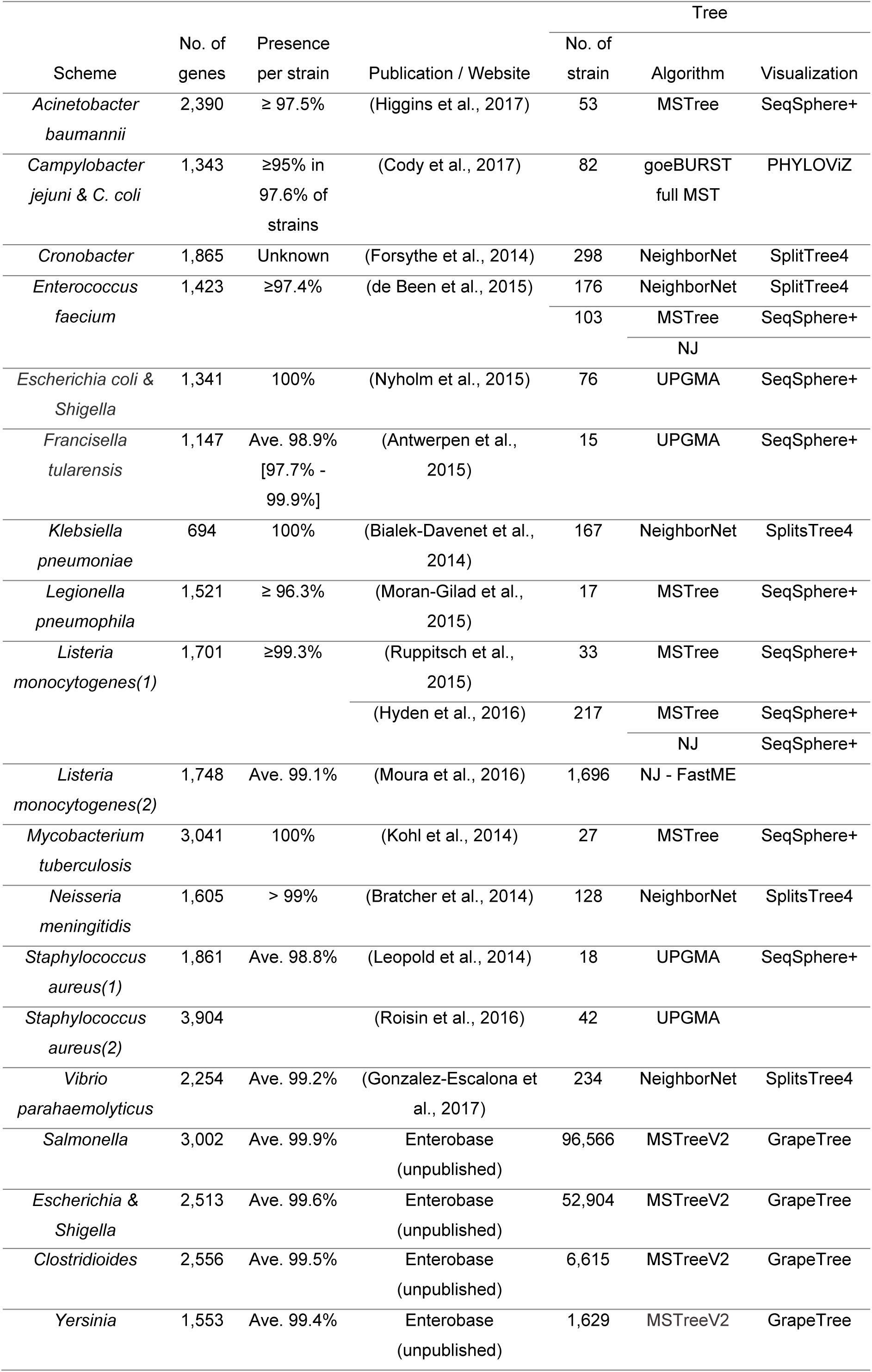
Analytical tools used in cgMLST/wgMLST studies (August, 2017)

### The big data conundrum

EnteroBase (https://enterobase.warwick.ac.uk) retrieves Illumina short reads from public repositories or reads that are uploaded by users for multiple genera, including *Salmonella* and *Escherichia*. It then assembles these reads into contigs, and assigns them to MLST STs at all levels of resolution from legacy MLST to wgMLST. EnteroBase also provides tools for investigating associations between metadata and genetically uniform populations. The development of EnteroBase began in 2014, at which time only few sets of short reads were available. In August 2017, it contained ~100,000 *Salmonella* genomes and >55,000 *Escherichia* genomes, and continues to grow rapidly. Such large numbers of assembled genomes plus their metadata facilitate comparisons of isolates from distinct geographical sources and over extended time scales. However, many existing methods for visualising genetic diversity in the form of dendrograms are not adequate to deal with these large datasets (Table 2). Even calculating a phylogram from so much data is challenging because phylogenetic tree inference algorithms, such as Neighbor-Joining (NJ), have a time complexity of O(n^3^) (Studier & Keppler, 1988), and *de novo*, sequence-based comparisons are not practical for large numbers of genomes. Exploring genomic datasets of this scale with current graphic visualisers is challenging because of the difficulty of appropriately representing both clusters of almost identical STs and deeper evolutionary relationships. For example, the default presentation by iTOL (Letunic & Bork, 2016) of 99,722 *Salmonella* spp. strains corresponding to 3,902 legacy MLST STs is not particularly intuitive (Fig. S1) even though iTOL is a very powerful tool that can show 100,000s of nodes.

**Table 2.** Selected tools for tree visualisation

|  | GrapeTree | Phyloviz 2.0 | SplitsTree4 | iTOL | EvoView | MicroReact | FigTree | Dendroscope | TreeDyn | TreeView |
| --- | --- | --- | --- | --- | --- | --- | --- | --- | --- | --- |
| Citation: | This study | (Nascimento et al., 2017) | (Huson & Bryant, 2006) | (Letunic & Bork, 2016) | (He et al., 2016) | (Argimon et al., 2016) | (Rambaut, 2016) | (Huson & Scornavacca, 2012) | (Chevenet et al., 2006) | (Page, 1996) |
| Front end | Enterobase | Phyloviz | BigsDB | iTOL | EvoView | MicroReact |  |  |  |  |
| Standalone version | ✓ | ✓ | ✓ |  |  |  | ✓ | ✓ | ✓ | ✓ |
| Rendering 100K nodes (time) | 47s* | Failed | Failed | 43s | Failed | Failed | 6s | 3s | Failed | Failed |
| Build 4K node tree (100K items) (time) | 15s* | Failed | Failed |  |  |  |  |  |  |  |
| Build 80K node tree (time) | 12hrs | Failed | Failed |  |  |  |  |  |  |  |
| MSTree | ✓ | ✓ |  |  |  |  |  |  |  |  |
| Database Integration | Enterobase |  | BigsDB |  |  | WGSA |  |  |  |  |
| Metadata Annotation | Size, Label, Colour | Size, Label | Label | Advanced | Advanced | Label, Colour | Size, Label, Colour | Label, Colour | Advanced |  |
| Metadata Operation | Sort, Filter, Edit | Filter |  | Sort, Filter |  |  | Filter |  |  |  |
| Inputs | Characters, Sequences, Nexus Tree | Characters, Sequences | Characters, Sequences, Nexus Tree | Nexus & XML Tree | Nexus & XML Tree | Nexus Tree | Nexus Tree | Nexus & XML Tree | Nexus Tree | Nexus Tree |
| Collapsing | Auto, Manual, Node size rescaling |  |  | Auto, Manual |  | Manual | Manual | Manual | Manual |  |
| Layout | Static, Dynamic & Free Moving | Dynamic & Free Moving | Static & Free Moving | Static | Static | Static | Static | Static | Static | Static |

Note: \*Times reported for GrapeTree to build and render trees are for the standalone version, and an additional 10-20s are required for the Enterobase version to load allelic profiles/metadata from the database. Static layout: computed before displayed; Dynamic layout: computed while displayed; Free Moving: Nodes are movable without changing the tree topology.

### The missing data problem

Even cgMLST genes are occasionally absent in individual assemblies (Table 1) due to deletions, or are not identified due to bioinformatic problems. The problem is further evident in wgMLST schemas, where the large majority of loci, since are part of an accessory genome, are lacking in individual genomes. Such uncalled alleles can result in large numbers of sets of almost identical STs, each of which is nevertheless a unique node in a phylogram because they differ in allelic content. As a result, >80,000 cgMLST STs have currently been called for *Salmonella* by EnteroBase, beyond the rendering capacities of many graphic visualizers. Minimum spanning trees have previously been used as an alternative to phylograms for representing clusters of related STs in 2-D space because they group isolates with identical STs within single nodes (Francisco et al., 2012; Nascimento et al., 2017). However, the branches and topologies based on cgMLST profiles in the minimum spanning tree (Fig. 1C) differ from the phylogeny calculated by NJ (Fig. 1B), or determined based on single nucleotide polymorphisms (SNPs) analysis (Fig. 1A), and are often distorted by missing data.

**Figure 1.**
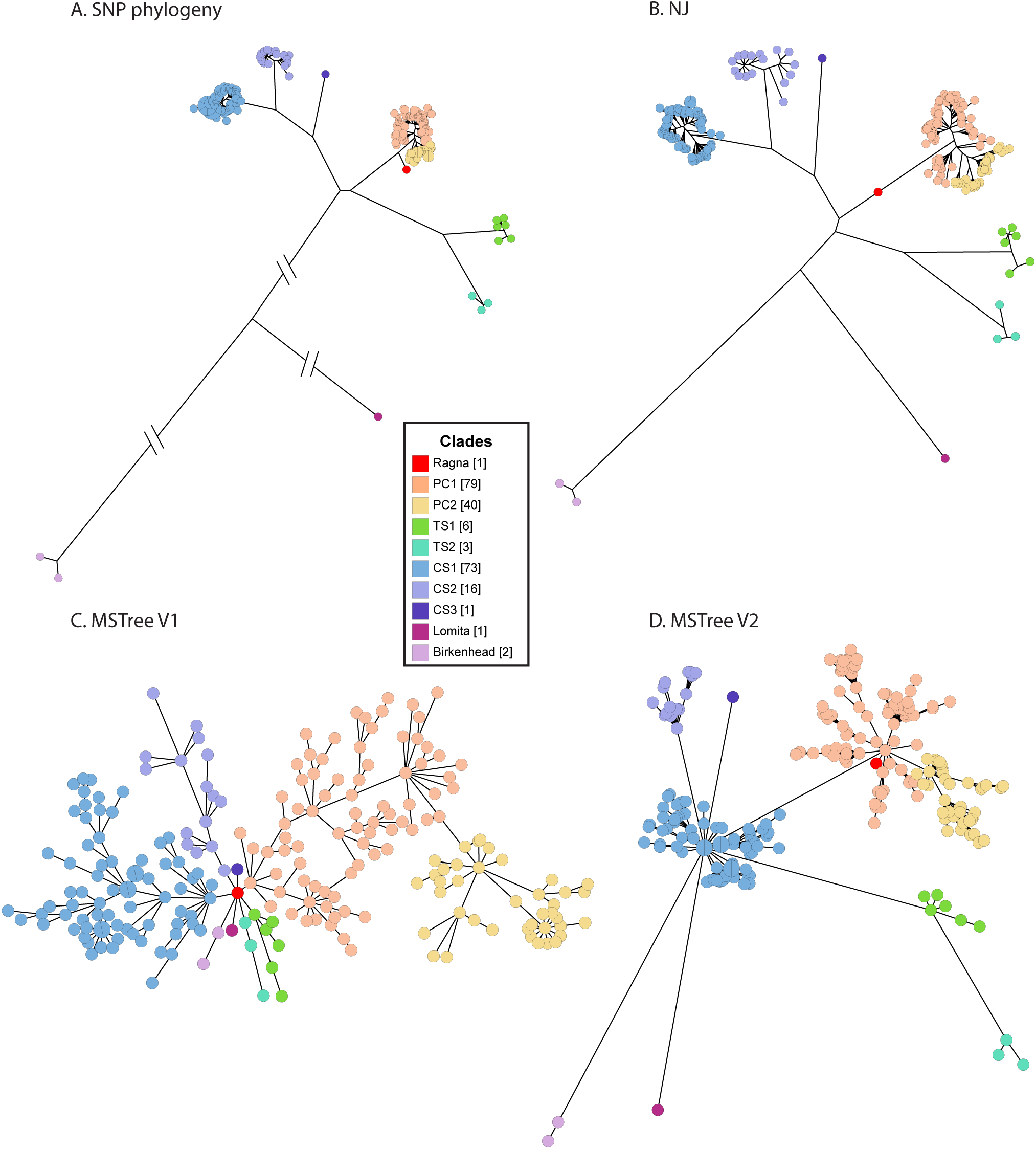
Comparisons of topologies produced by four algorithms from 49,610 non-recombinant, core genomic SNPs (A) or 3,002 loci in the cgMLST V2 scheme (B-D) from 219 genomes within the Para C Lineage (Zhou et al., 2017). Trees were reconstructed based on Maximum Likelihood (A), Neighbour-Joining (B), MSTree (C) or MSTreeV2. Nodes are colour-coded by Clade (key).The 219 Para C Lineage genomes were supplemented by an ancient genome (Ragna; red) reconstructed from an 800-yr old skeleton (Zhou et al., 2017) plus two related but distinct genomes of serovar Birkenhead, which act as an outgroup. Alleles were only called for 215 of the 3,002 cgMLST loci from Ragna (<10%) and the remainder represent missing data due to the fragmentation in ancient DNA. Despite the high levels of missing data, the neighbour-joining method (B) reconstructed a tree with similar topology to that of the SNP phylogeny (A), except that the PC Clades branch from the main branch in B and Ragna is located directly on the branch leading to the PC Clades. All Clades that are distinct within the SNP phylogeny also form distinct clusters according to the MSTreeV2 tree (D), but not in the minimum spanning tree (C). Instead, the minimum spanning tree topology radiates from Ragna, which has the smallest number of allelic differences to all other genotypes simply because most alleles are scored as missing data. Interactive versions of the figures are publicly available to registered EnteroBase users via links in a public workspace https://goo.gl/Phrm4f.

Here we present GrapeTree, a web application that reconstructs and visualises intricate phylogenetic trees together with detailed metadata. GrapeTree supports facile manual manipulations of both tree layout and metadata, or by setting threshold values, and is fully interactive. GrapeTree is available in a standalone (SA) version (Fig S2) which handles pre-calculated trees plus metadata in text form or fully integrated into Enterobase (EB) (Fig. 1D), thus leveraging information from 100,000s of bacterial genomes and their associated metadata.

## GrapeTree: visualization

### Large datasets

GrapeTree can handle large datasets, such as the relationships of 99,722 *Salmonella* genomes that were assigned by EnteroBase to 3,902 legacy MLST STs (Fig. 2). Previous analyses of a 24-fold smaller sample of *Salmonella* isolates (Achtman et al., 2012) had clustered such STs in eBurstGroups (eBGs), which were predominantly uniform for serological surface properties (serovar). Colour coding by the serovar predictions calculated by EnteroBase provides visual confirmation of these observations (Fig. 2). This graphic presentation was completed in under 1.5 minutes (Table 2).

**Figure 2.**
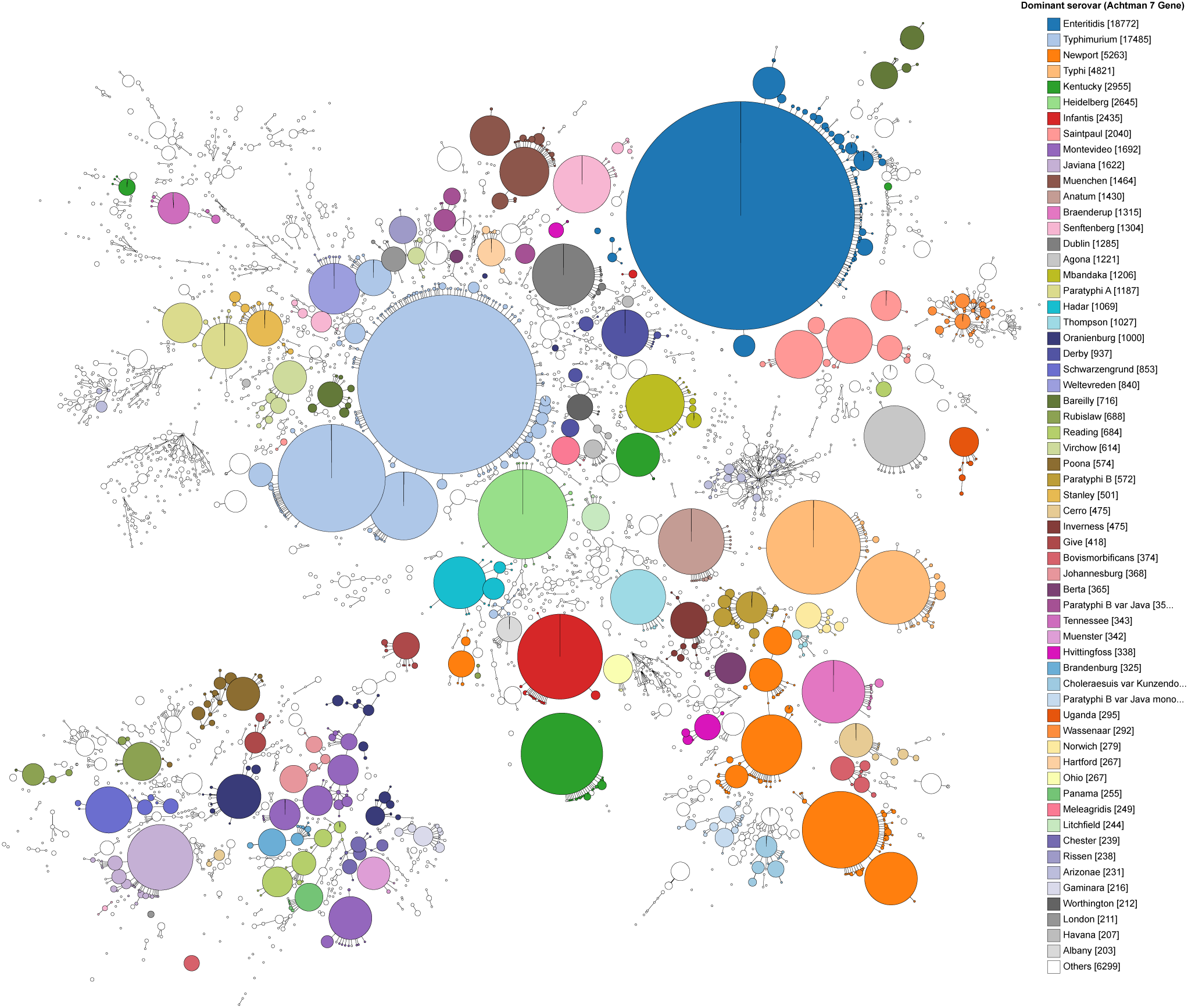
GrapeTree (EB) clustering with MSTreeV2 of 3,902 legacy MLST STs from 99,722 genomic assemblies in *Salmonella* EnteroBase. Each node corresponds to a single ST, with diameter scaled to the number of assemblies, and was colour-coded according to the dominant serovar of the corresponding eBURST group (Achtman et al., 2012). Colours associated with the 60 most prevalent serovars are indicated in the key (right). Edges indicate differences between STs of 1-2 of the 7 loci. Time needed for calculation and rendering: 1.5 min. An interactive version of the figure is publicly available to registered EnteroBase users at http://enterobase.warwick.ac.uk/ms_tree?tree_id=6168.

### Flexibility

GrapeTree (SA) accepts tables of character data (allelic profiles or SNPs), pre-calculated tree files in standard formats (Newick or NEXUS), and comma-delimited text for metadata (Fig. S2). Data can be input into SA by dragging and dropping files from a user’s local workstation, or from online sources. The SA backend module calculates trees from character data whereas pre-calculated tree files are displayed without further modification other than rendering. To illustrate these features, Fig. 3 shows a phylogenetic tree of 1,610 Ebola genomes from the 2013-2016 Ebola epidemic in West Africa (Dudas et al., 2017) which was downloaded together with associated metadata from MicroReact (Argimon et al., 2016).

**Figure 3.**
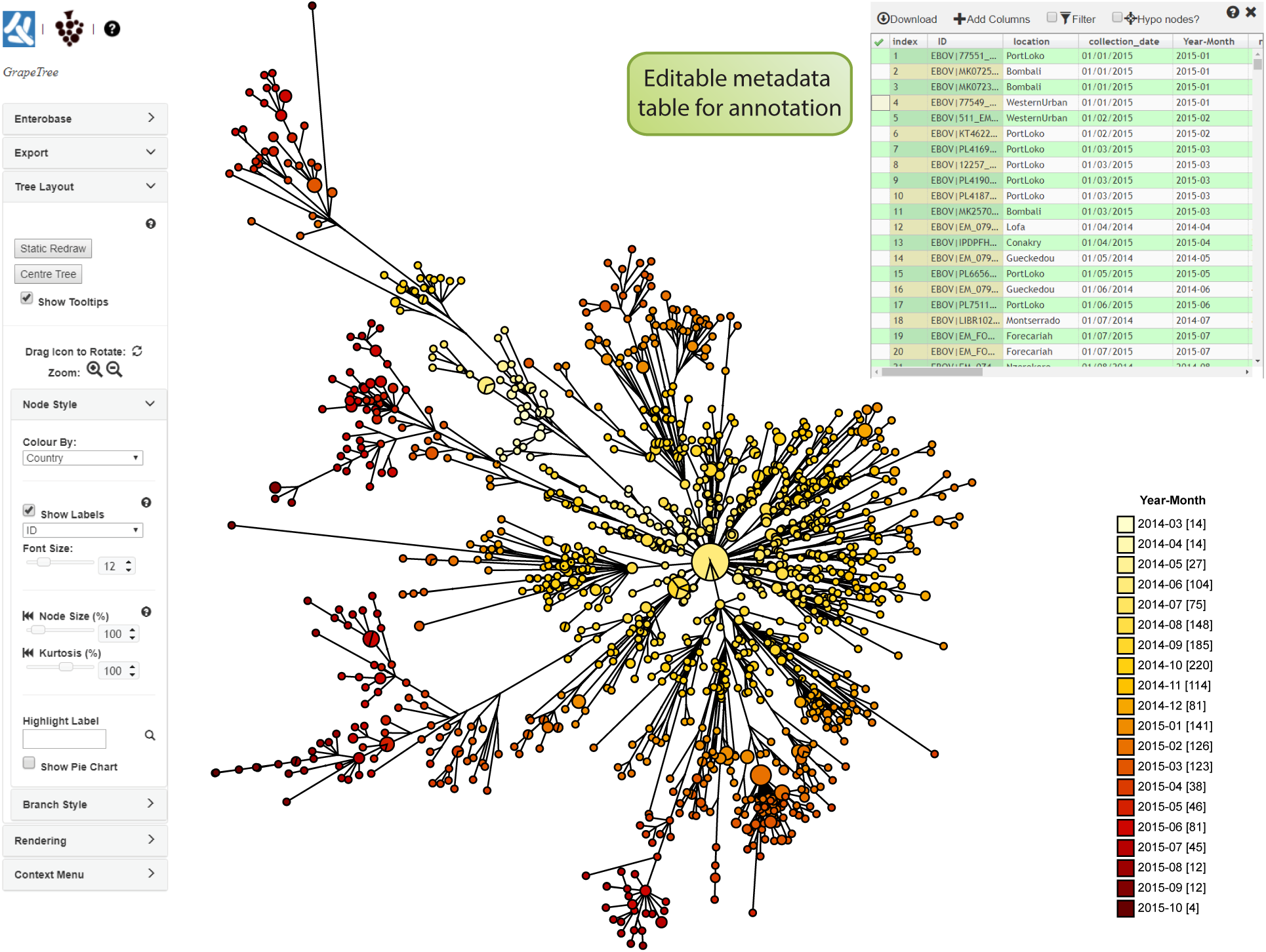
GrapeTree (SA) interface exemplified with a pre-calculated Newick tree based on 1,610 Ebola genomes from the West African epidemic of 2013-2016. The tree and metadata including a column designated “collection_date” were retrieved from microreact.org (https://microreact.org/project/west-african-ebola-epidemic). A new data column (Year-Month, upper right) was added to the metadata panel which contained the year and month information from “collection_date”, and this column was used to colour-code the visualisation as a temporal gradient (key, lower right). Branches spanning <0.22 substitutions per site were collapsed for clarity. The data indicate progressive radiation from a central source, consistent with published findings (Dudas et al., 2017). An interactive version of this figure and metadata can be found at https://goo.gl/iKJRny. The detailed functionality of GrapeTree is summarised in Fig. S2.

### Rendering

Complex trees are difficult to visualise with clarity. In order to address this issue, GrapeTree implements a modified version of the Equal-Daylight algorithm (Felsenstein, 2004a) to initialize node positioning (Fig. S3). The Equal-Daylight algorithm attempts to prevent overlapping child nodes by splitting the tree layout task into a series of sequential node layout tasks (Fig. S3). Our modification (Appendix 5) provides a solution to this task in linear time complexity. The resulting layout can be further adjusted dynamically on the entire tree, or on selected sub-trees, using the force-directed algorithm (Dwyer, 2009) in the JS D3 library (Appendix 4). Users can also manually adjust the branches of a tree to a preferred layout by rotating selected nodes and branches.

Many other visual aspects of a GrapeTree graph are also customizable by users (Fig. S2). In particular, simplifying complex trees with numerous nodes can be addressed by manually collapsing subsets of related nodes, or by setting a global threshold of differences below which all related nodes are collapsed (Appendix 6). The relationship between node size and number of entries can be adjusted in absolute terms, or by adjusting kurtosis. Branches can be cropped or hidden if their length is below a given threshold, e.g. in Fig. 3, branches of <0.22 substitutions per site were collapsed for clarity. The display of branch and/or node labels can be toggled.

### Metadata

GrapeTree implements a metadata panel based on SlickGrid (https://github.com/6pac/SlickGrid) that allows users to view and modify metadata that are associated with the individual entries (Fig. 3, top right). Any column can be used to colour code and/or label tree nodes. For example, in order to obtain an attractive presentation, the temporal gradient in Fig. 3 was implemented by creating a new metadata column (Year-Month) from data which had been downloaded in a different format, and specifying that colour coding was in gradient format. Similarly, although colour codes are assigned automatically to individual keys, those codes can be changed manually and the number of keys can be adjusted. In the metadata panel, metadata columns can be sorted and/or filtered at will to focus on entries of interest, and those entries that are selected in the metadata panel are immediately highlighted in the tree.

### Outputs

Both versions of GrapeTree support the export of the current state of the browser window, including the tree layout and all metadata, as a JSON file for use in future GrapeTree sessions. The figure and its underlying tree and metadata can be independently exported for manipulation with other software in Scalar Vector Graphics (SVG) and Newick tree formats, respectively. Local metadata (SA) can be saved in tab-delimited text format and metadata that are modified in EB GrapeTree can be uploaded to EnteroBase. Furthermore, nodes that were selected within EB can be loaded into an EnteroBase workspace for further processing.

## GrapeTree: algorithms

### NJ and MSTree

Multiple distance-based approaches are available for calculating trees from limited numbers of genomes (Table 1), and EnteroBase implements three of them. FastME V2 (Lefort et al., 2015) is used to calculate NJ trees from either categorical MLST data at all levels of resolution, or SNPs relative to a user-selected reference genome. However, FastME is not capable of handling the enormous numbers of genomes already present in EnteroBase (Fig. S4). GrapeTree therefore also implements a classical minimum spanning tree approach (MSTree) for MLST data, which is based on the Kruskal algorithm (Kruskal, 1956) with tie-breaking between multiple co-optimal branches according to the principles of eBURST (Feil et al., 2004) and its successor, goeBURST (Francisco et al., 2009). Fig. 1 presents the results from all three approaches tested on a relatively uniform group of 222 *Salmonella* genomes from the Para C Lineage, including one ancient genome which contained large amounts of missing data (Zhou et al., 2017). The cgMLST analysis with NJ (Fig. 1B) yielded very similar topologies to a maximum-likelihood tree from non-recombinant SNPs (Fig. 1A) with one exception: the SNP analysis correctly placed the 800 year old ancient DNA (red node) on a side branch relative to the modern members of serovar Paratyphi C whereas that node was inaccurately placed on the main branch in the cgMLST tree.

The handling of cgMLST data was poorer with the classical MSTree algorithm than with NJ (Fig. 1C). The clustering of closely-related modern genomes was not as clearly defined, and the topological branching order and branch lengths were drastically different. This is in accord with our general experience that the classical minimum spanning tree algorithm erroneously places nodes with extensive missing data in central positions, such as the ancient DNA in this analysis, and generally draws faulty topologies when confronted with missing data. We therefore developed MSTreeV2, a highly improved algorithm (Fig. S5A) for inferring genetic relationships from 10,000s of allelic profiles in quadratic time complexity, including missing data (Fig. 1D). EnteroBase offers MSTreeV2 as its default tree-building algorithm for GrapeTree visualization but continues to offer the others as alternatives.

### MSTreeV2

The classical minimum spanning tree is based on non-directional distances, which does not penalise nodes containing missing data. However, cgMLST analyses need a directional metric because most missing data in genomic sequences arises from technical errors related to sequencing or assembly, rather than biological changes such as deletions and insertions. MSTreeV2 calculates a directed minimum spanning tree (dMST; also called a minimum spanning arborescence) based on asymmetric Hamming-like distances in which the directionality is from more complete to less complete profiles (Fig. S5B). In addition, eBURST-based minimum spanning trees try to identify ‘founders’ of clustered nodes, and use the simple metrics defined by the eBURST/goeBURST heuristic as tiebreakers between co-optimal edges. However, minimum spanning trees lack hypothetical nodes, which can result in distorted topologies when founder nodes do not exist in cgMLST data from modern genomes. Instead of searching for a possibly non-existing founder node, MSTreeV2 identifies “centroid” genotypes with the lowest harmonic mean allelic distance to all other genotypes in a population, thereby preferentially weighting smaller allelic distances between variant STs. Finally, standard and naive minimum spanning trees implementations do not use evolutionary principles in choosing among co-optimal edges. Therefore, MSTreeV2 subjects the final tree to local branch recrafting depending on the maximum likelihood fit to two distinct phylogenetic models (Fig. 5C-E). The mathematical principles underlying these processes are presented in Appendices 1-3.

## Comparative analyses of simulated data

SimBac (Brown et al., 2016) was used to simulate the coalescence of 40 genomes with 100 replicates for each of 24 different substitution rates at a constant population size and no homologous recombination. Allelic profiles for 2,000 loci were then called for the simulated sequences (Appendix 7). The effect of missing values was also evaluated by removing randomly selected values from the allelic profiles for substitution rate 0.00005.

We inferred trees for each replicate with classical NJ (Felsenstein, 2004b). We also inferred MSTrees using command line versions of goeBURST (Francisco et al., 2009), MSTreeV2, and dMST (the intermediate result of MSTreeV2 immediately preceding local branch recrafting). For simulations with missing values, we compared goeBURST with missing data treated as a common additional allele (goeBURST[a]) and with missing data being ignoring in pairwise comparisons (goeBURST[i]). The algorithms were measured for both precision (frequency of true positives) and sensitivity (inverse frequency of false positives) for reconstructing quartet splits (Strimmer & von Haeseler, 1996) against the known history of evolutionary changes in the simulated data. The simulated data arose via binary splits and does not include any star-like quartets. Minimum spanning trees tend to create star-like quartets, which are unresolved, and were also scored as false negatives.

The results in Fig. 4A show that both MSTreeV2 and NJ performed very well in regards to precision with complete allelic profiles, much better than both variants of goeBURST, which is increasingly less precise as the allelic distance increases between allelic profiles. Importantly, the high precision of MSTreeV2 also applies to missing data (Fig. 4B). NJ is much better than any of the minimal spanning methods in regard to sensitivity, arguably because it accounts for hypothetical nodes (Appendix 7). However, much larger datasets can be visualised by minimum spanning trees than by NJ (Fig. S4).

**Figure 4.**
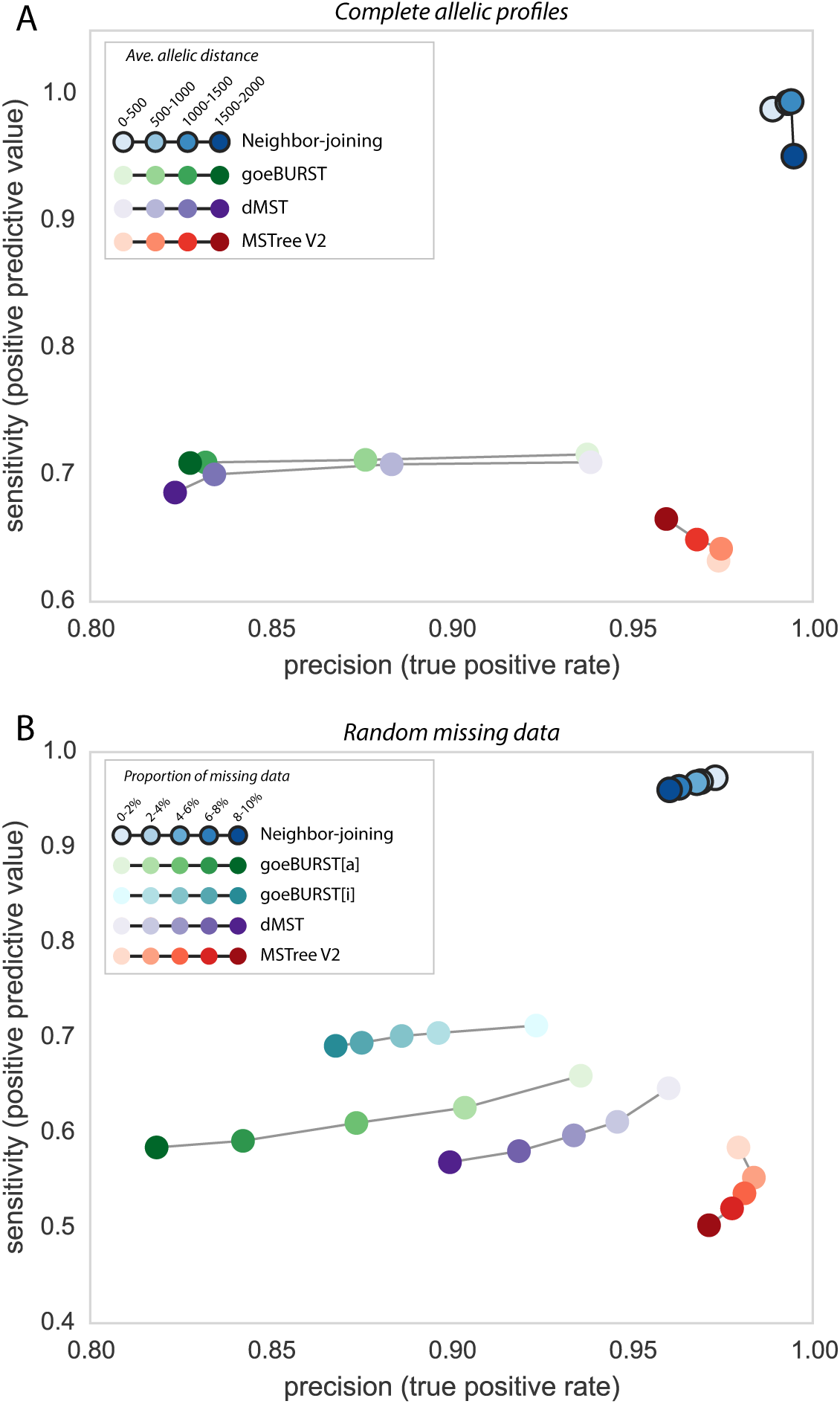
Sensitivity and precision of trees calculated from simulated allelic data by multiple algorithms. Trees were calculated for 100 replicas from each of 24 simulated phylogenies which differed in substitution rates between 0.00001 and 0.07. A) Sensitivity *vs* precision after quartet analysis of branches calculated in the absence of missing data by NJ, the goeBURST algorithm used in MSTree, the dMST intermediate stage prior to local branch recrafting, and the full MSTreeV2 algorithm. The results with each algorithm are summarised by four values after binning by allelic distances within the quartets. B) Sensitivity *vs* precision after quartet analysis of branches calculated with different levels of random missing data for substitution rate 0.00005. goeBURST was forced to treat missing values as additional alleles by encoding them as 0 (goeBURST[a]) or to ignore them by encoding them as ‘-’ (goeBURST[i]; default in MSTree). The results with each algorithm are summarised by five average values after binning by the proportion of missing data.

The evaluation also showed that precision was similar between dMST and goeBURST with complete allelic profiles, confirming the equivalence of both approaches in the absence of missing data. In contrast the precision of dMST was higher with missing data than for either goeBURST algorithm, which demonstrates the benefits of the directed MST approach adopted in the intermediate step of MSTreeV2. Precision with MSTreeV2 was even higher than with dMST, indicating the importance of the local branch recrafting in MSTreeV2 for its accuracy of calls. The trade-off is that the high precision is accompanied by a slightly lower sensitivity than is true of classical minimum spanning trees. The analyses in Fig. S6 indicate that the slightly lower sensitivity of MSTreeV2 in comparison to classical MSTree algorithms is because it does not attempt to resolve any topologies for unbalanced quartets (Appendix 7).

## Conclusion

Core genome MultiLocus Sequence Typing provides a feasible approach for providing public access to 100,000s of bacterial genotypes at the genomic level. Access to such databases will facilitate international collaboration and support the global surveillance of bacterial pathogens. The current bottleneck is that of real-time graphical visualisation of the relationships between such large datasets. GrapeTree satisfies that need, and allows users to explore the population structure and phenotypic properties of large numbers of genomes in a web browser with fine-grained resolution. GrapeTree is available as a graphical frontend to EnteroBase, providing access to cgMLST schemes of multiple bacterial pathogens as well as in a standalone version that allows exploration of user-define trees. GrapeTree empowers the public exploitation of genomic data by non-bioinformaticians and can close the current gap between epidemiology and genomics.

## Acknowledgements

EnteroBase (BBSRC BB/L020319/1) was developed by N-F.A., M.J.S. and Z.Z. (equal contributions) under guidance by M.A. Additional grant support was from the Wellcome Trust (202792/Z/16/Z). We thank Philipe Lemey and Joseph Healey for beta testing and feedback.

## Author Contributions

ZZ and MJS conceived the ideas and designed methodology and functionality; MJS, ZZ, NA and MA designed look and feel of the GrapeTree GUI; ZZ and NL performed simulations and analysed the data; APF and CV contributed to the formalization, correctness and analysis of algorithms; APF implemented the command line version of goeBURST; JAC reviewed concepts and implementation of MST and associated algorithms. MA led the writing of the manuscript. All authors contributed critically to the drafts and gave final approval for publication.

## Data Accessibility

Interactive versions of Figs. 1–3 can be found at https://goo.gl/Phrm4f, http://enterobase.warwick.ac.uk/ms_tree?tree_id=6168 and https://goo.gl/iKJRny. Source code is available at https://github.com/martinSergeant/EnteroMSTree/ and precompiled binaries, online documentation and a live demo are available at https://bitbucket.org/enterobase/enterobase-web/wiki/GrapeTree.

---

**Figure S1. Visualisation in iTOL of data from Fig. 2.** The tree in Fig. 2 based on 3,902 MLST STs from 99,722 *Salmonella* isolates was used to generate a Newick file in GrapeTree. That Newick file was used as input to iTOL (Letunic & Bork, 2016) and the tips were colour-coded according to the 60 most prevalent serovars as in Fig. 2 (key). An interactive version of this tree can be found in http://itol.embl.de/tree/13720512318460241506332484.

**Figure S2. Features implemented in GrapeTree.** All features are common to the Standalone and EnteroBase versions, except where their specificity to one version is indicated by SA or EB.

**Figure S3. Node layout algorithm in GrapeTree.** The node layout depends on the calculation of descendent circular sectors (DCS) for each node. Node *n* has two children nodes, *c*_*1*_ and *c*_*2*_. All the descendants (dotted circles) of *c*_*1*_ are encompassed within DCS(*c*_*1*_) (cyan), whereas *c*_*2*_ has no descendants. Node *c*_*2*_ is larger than either *n* or *c*_*1*_ because it contains multiple entries. A) Initial calculation. B) Child’s circular sector CSS(*c*_*1*_) (green, left) is drawn to include both *c*_*1*_ and all its descendants. C) CSS(*c*_*2*_) (green, right) is drawn to include large node *c*_*2*_. D) DCS(*n*) is drawn as a summarized circular sector (dotted lines) that includes both CSS(*c*_*1*_) and CCS(*c*_*2*_) plus a separating arc *s* (red) between them.

**Figure S4. Time profiling of analytic tools with simulated data.** The time to completion was determined on 10 replicates each of simulated data for 1,000-10,000 distinct STs with two variants of NJ (Phylip (Felsenstein, 2004b); FastME (Lefort et al., 2015)), two variants of a minimal spanning tree (goeBURST (Nascimento et al., 2017); MSTreeV2) and NeighborNet from Splitstree4 (Huson & Bryant, 2006; Huson & Bryant, 2017). In all cases, pre-computed distance matrices were used as inputs for the algorithms except for goeBURST, which only accepts allelic profiles as inputs. Times are the minimum of three independent runs for each replica and algorithm. Computations for which results are not shown were terminated with completion at 4 hours.

**Figure S5. Workflow of the MSTreeV2 algorithm.** A) Sequential steps. See Appendix for greater details. B) Calculation of asymmetric Hamming-like distance for all possible combinations of allelic values in the calculation of the edge distance *d(u→v)*. The genetic distance for each locus within an ST profile is 0 when the locus contains the same allele in *u* and *v*, and 1 when that locus contains distinct alleles. It is 1 when *v* contains an existing allele at that locus but the allele is missing in *u*, and is otherwise 0. C) Detailed workflow for the local branch recrafting of edge *u→v*. D) Model A: nodes *u* and *w* are contemporary sisters that diverged from a hypothetical common ancestor. E) Model B: node *w* is the direct ancestor of *u*. The probabilities of each model and the branch parameters *l*_*A*_, *k*_*A*_, *l*_*B*_ and *k*_*B*_ are calculated using equations 1-2 from Appendix 3.

**Figure S6. Sensitivity *vs* precision for balanced and unbalanced quartets.** A) Cartoon of balanced quartet. B) Cartoon of unbalanced quartet. C-D) Quartets are binned according to their average allelic distance as in Fig. 4. C) Performance for balanced quartets. All algorithms perform well. D) Performance for unbalances quartets. NJ resolves unbalanced quartets accurately, but the sensitivity is <0.6 for all minimum spanning tree algorithms. In addition, precision is low for goeBURST and dMST but remains quite high with MSTreeV2 because local branch recrafting removes most of the inaccurate splits.

**Figure S7. A cartoonised example for the branch recrafting.** Green lines in (B, D, F) are the branches involved in the model comparisons, in which solid lines are the previously chosen branches and dotted branches are the proposed branches. Red lines in (C, E, G) show the outputs of the model comparisons, in which Solid lines are the most probable branches and the dotted lines are the less possible ones. (A) The shortest branch *(E→A)* inferred by Edmond’s algorithm that connects two trees *t(E)* and *t(A)*. (B) *(E→A)* is compared with branch *(F→A)*, where node *F* has the lowest harmonic average distance to other nodes. (C) *(F→A)* has higher probability. (D) *(F→A)* is compared with all the nodes that are directly connected with F. (E) *(F→A)* is still the most probable branch. (F-G) The same process is done for tree *t(A)*, which results to the most probably branch *(F→B)*.

## References

Achtman, M., Wain, J., Weill, F.-X., et al. (2012) Multilocus sequence typing as a replacement for serotyping in Salmonella enterica. PLoS Pathogens, 8, e1002776.

Antwerpen, M.H., Prior, K., Mellmann, A., et al. (2015) Rapid high resolution genotyping of *Francisella tularensis* by whole genome sequence comparison of annotated genes ("MLST+"). PLoS.One., 10, e0123298.

Argimon, S., Abudahab, K., Goater, R.J., et al. (2016) Microreact: visualizing and sharing data for genomic epidemiology and phylogeography. Microb.Genom., 2, e000093.

Bialek-Davenet, S., Criscuolo, A., Ailloud, F., et al. (2014) Genomic definition of hypervirulent and multidrug-resistant *Klebsiella pneumoniae* clonal groups. Emerg.Infect.Dis., 20, 1812–1820.

Bratcher, H.B., Corton, C., Jolley, K.A., Parkhill, J. & Maiden, M.C. (2014) A gene-by-gene population genomics platform: *de novo* assembly, annotation and genealogical analysis of 108 representative *Neisseria meningitidis* genomes. BMC.Genomics, 15, 1138.

Brown, T., Didelot, X., Wilson, D.J. & De, M.N. (2016) SimBac: simulation of whole bacterial genomes with homologous recombination. Microb.Genom., 2.

Chevenet, F., Brun, C., Banuls, A.L., Jacq, B. & Christen, R. (2006) TreeDyn: towards dynamic graphics and annotations for analyses of trees. BMC.Bioinformatics., 7, 439.

Cody, A.J., Bray, J.E., Jolley, K.A., McCarthy, N.D. & Maiden, M.C.J. (2017) Core genome multilocus sequence typing scheme for stable, comparative analyses of *Campylobacter jejuni* and *C. coli* human disease isolates. J.Clin.Microbiol., 55, 2086–2097.

de Been, M., Pinholt, M., Top, J., et al. (2015) Core genome multilocus sequence typing scheme for high-resolution typing of *Enterococcus faecium*. J.Clin.Microbiol., 53, 3788–3797.

Dudas, G., Carvalho, L.M., Bedford, T., et al. (2017) Virus genomes reveal factors that spread and sustained the Ebola epidemic. Nature (London), 544, 309–315.

Dwyer, T. (2009) Scalable, versatile and simple constrained graph layout. Eurographics, 28.

Feil, E.J., Li, B.C., Aanensen, D.M., Hanage, W.P. & Spratt, B.G. (2004) eBURST: Inferring patterns of evolutionary descent among clusters of related bacterial genotypes from Multilocus Sequence Typing data. Journal of Bacteriology, 186, 1518–1530.

Felsenstein, J. (2004a) Drawing trees. Inferring phylogenies. pp. 573–584. Sinauer Associates, Inc., Sunderland, Massachusetts, US.

Felsenstein, J. (2004b) PHYLIP (Phylogeny Inference Package) version 3.6b. Available at: http://evolution.genetics.washington.edu/phylip.html.

Forsythe, S.J., Dickins, B. & Jolley, K.A. (2014) *Cronobacter*, the emergent bacterial pathogen *Enterobacter sakazakii* comes of age; MLST and whole genome sequence analysis. BMC.Genomics, 15, 1121.

Francisco, A.P., Bugalho, M., Ramirez, M. & Carrico, J.A. (2009) Global optimal eBURST analysis of multilocus typing data using a graphic matroid approach. BMC Bioinformatics., 10, 152.

Francisco, A.P., Vaz, C., Monteiro, P.T., et al. (2012) PHYLOViZ: phylogenetic inference and data visualization for sequence based typing methods. BMC Bioinformatics, 13, 87.

Gonzalez-Escalona, N., Jolley, K.A., Reed, E. & Martinez-Urtaza, J. (2017) Defining a core genome multilocus sequence typing scheme for the global epidemiology of *Vibrio parahaemolyticus*. J.Clin.Microbiol., 55, 1682–1697.

He, Z., Zhang, H., Gao, S., et al. (2016) Evolview v2: an online visualization and management tool for customized and annotated phylogenetic trees. Nucleic Acids Res., 44, W236–W241.

Higgins, P.G., Prior, K., Harmsen, D. & Seifert, H. (2017) Development and evaluation of a core genome multilocus typing scheme for whole-genome sequence-based typing of *Acinetobacter baumannii*. PLoS.One., 12, e0179228.

Huson, D.H. & Bryant, D. (2006) Application of phylogenetic networks in evolutionary studies. Molecular Biology and Evolution, 23, 254–267.

Huson, D. H. & Bryant, D. (2017) SplitsTree4. Available at: http://www.splitstree.org/.

Huson, D.H. & Scornavacca, C. (2012) Dendroscope 3: an interactive tool for rooted phylogenetic trees and networks. Syst.Biol, 61, 1061–1067.

Hyden, P., Pietzka, A., Lennkh, A., et al. (2016) Whole genome sequence-based serogrouping of *Listeria monocytogenes* isolates. J.Biotechnol., 235, 181–186.

Jolley, K.A. & Maiden, M.C. (2014) Using multilocus sequence typing to study bacterial variation: prospects in the genomic era. Future.Microbiol, 9, 623–630.

Kohl, T.A., Diel, R., Harmsen, D., et al. (2014) Whole-genome-based *Mycobacterium tuberculosis* surveillance: a standardized, portable, and expandable approach. J.Clin.Microbiol., 52, 2479–2486.

Kruskal, J.B. (1956) On the shortest spanning subtree of a graph and the traveling salesman problem. Proceedings of the American Mathematical Society, 7, 48–50.

Lefort, V., Desper, R. & Gascuel, O. (2015) FastME 2.0: A Comprehensive, Accurate, and Fast Distance-Based Phylogeny Inference Program. Mol.Biol.Evol., 32, 2798–2800.

Leopold, S.R., Goering, R.V., Witten, A., Harmsen, D. & Mellmann, A. (2014) Bacterial whole-genome sequencing revisited: portable, scalable, and standardized analysis for typing and detection of virulence and antibiotic resistance genes. J.Clin.Microbiol., 52, 2365–2370.

Letunic, I. & Bork, P. (2016) Interactive tree of life (iTOL) v3: an online tool for the display and annotation of phylogenetic and other trees. Nucleic Acids Research, 44, W242–W245.

Maiden, M.C., van Rensburg, M.J., Bray, J.E., et al. (2013) MLST revisited: the gene-by-gene approach to bacterial genomics. Nature Reviews Microbiology, 11, 728–736.

Maiden, M.C.J., Bygraves, J.A., Feil, E., et al. (1998) Multilocus sequence typing: A portable approach to the identification of clones within populations of pathogenic microorganisms. Proceedings of the National Academy of Sciencs of the United States of America, 95, 3140–3145.

Moran-Gilad, J., Prior, K., Yakunin, E., et al. (2015) Design and application of a core genome multilocus sequence typing scheme for investigation of Legionnaires’ disease incidents. Euro.Surveill, 20.

Moura, A., Criscuolo, A., Pouseele, H., et al. (2016) Whole genome-based population biology and epidemiological surveillance of *Listeria monocytogenes*. Nat Microbiol, 2, 16185.

Nadon, C., Van, W., I, Gerner-Smidt, P., et al. (2017) PulseNet International: Vision for the implementation of whole genome sequencing (WGS) for global food-borne disease surveillance. Euro.Surveill, 22.

Nascimento, M., Sousa, A., Ramirez, M., et al. (2017) PHYLOViZ 2.0: providing scalable data integration and visualization for multiple phylogenetic inference methods. Bioinformatics., 33, 128–129.

Nyholm, O., Halkilahti, J., Wiklund, G., et al. (2015) Comparative genomics and characterization of hybrid shigatoxigenic and enterotoxigenic *Escherichia coli* (STEC/ETEC) strains. PLoS.One., 10, e0135936.

Page, R.D. (1996) TreeView: an application to display phylogenetic trees on personal computers. Comput.Appl.Biosci., 12, 357–358.

Rambaut, A. (2016) FigTree v.1.4.3. Available at: http://tree.bio.ed.ac.uk/software/figtree/.

Roisin, S., Gaudin, C., de Mendonca, R., et al. (2016) Pan-genome multilocus sequence typing and outbreak-specific reference-based single nucleotide polymorphism analysis to resolve two concurrent *Staphylococcus aureus* outbreaks in neonatal services. Clin.Microbiol.Infect., 22, 520–526.

Ruppitsch, W., Pietzka, A., Prior, K., et al. (2015) Defining and evaluating a core genome multilocus sequence typing scheme for whole-genome sequence-based typing of *Listeria monocytogenes*. J.Clin.Microbiol., 53, 2869–2876.

Strimmer, K. & von Haeseler, A. (1996) Quartet puzzling: a quartet maximum-likelihood method for reconstructing tree topologies. Molecular Biology and Evolution, 13, 964–969.

Studier, J.A. & Keppler, K.J. (1988) A note on the neighbor-joining algorithm of Saitou and Nei. Molecular Biology and Evolution, 5, 729–731.

Zhou, Z., Lundstrøm, I., Tran--Dien, A., et al. (2017) Millenia of genomic stability within the invasive Para C Lineage of Salmonella enterica. BioRxiv.

